# Training and transfer effects on motor skill learning: different encoding processes that lead to mechanism-specific neural codes

**DOI:** 10.1101/2025.07.03.662838

**Authors:** Theo Marins, Frederico Augusto Casarsa de Azevedo, Fernanda Tovar-Moll, Guilherme Wood

## Abstract

Motor learning typically emerges from repetitive practice but can also arise through transfer or generalization, when improvements extend to untrained actions. However, it remains unclear whether training-based learning (training effect) and transfer/generalization-based learning (transfer effect) rely on shared neural mechanisms or whether they represent distinct processes supported by different cortical architectures. Here, we employed between-subject multivariate decoding of hyperaligned functional magnetic resonance imaging data to characterize the universal neural codes underlying the training and transfer effects and their resultant motor engrams in the sensorimotor cortex. We found that both learning processes are encoded bilaterally but shifted to the right (ipsilateral) hemisphere, with the training effect showing higher decoding accuracy than the transfer effect. The bilateral central sulci showed a clear mechanism-specificity towards the training effect, in contrast to the rest of the sensorimotor cortex. Interestingly, these learning processes consolidate two distinct motor engrams. Remarkably, the information structure of these processes is shared among individual brains, and the between-subject classification provided limited results when performed with standard, anatomical alignment. Our findings support the notion that motor skill learning is hierarchically organized in the sensorimotor cortex, and extend current models by revealing that training and transfer encode distinct and mechanism-dependent high-dimensional representations embedded in a neural code that is shared across brains.

## Introduction

Motor learning often requires repetitive training, but it is also achieved through mechanisms of transfer of learning or generalization, in which improvements extend to untrained conditions (1). In motor training of finger-tapping sequences, this is detected as sequence-independent learning and suggests that both mechanisms of learning share similar neural resources (2, 3). Moreover, neuroimaging studies have reported bilateral activation of M1/S1, cerebellum, and parietal regions during both training and transfer phases (2–4), but also have shown differential sensitivity to age (5) and task manipulations (6, 7), suggesting that the two learning mechanisms are partially distinct at the neural level. Altogether, previous studies support the hierarchical view of motor skill learning (8), which predicts that the repetitive selection of small movement elements leads to the formation of reusable representations of fragments of sequential movements, or chunks. In this way, the low-level representation of sequential movements is stably encoded in the primary motor (M1) and somatosensory cortices (S1), while higher-level sequence representation relies on secondary sensorimotor areas (8–11).

In the present study, we investigated the neural codes of training-based and transfer-based learning and described the motor engrams resulting from these learning processes. We used hyperalignment (12) to derive a common representational space that captures the basis functions that are shared among individuals. Hyperalignment leads to above-chance accuracy in decoding models applied to the visual system (13–17) by eliminating idiosyncratic, noisy, and task-unrelated neural patterns. A recent study has shown that hyperalignment reveals a common neural representation of finger-tapping movements (18). We show that the neural representations of both learning mechanisms are shared across individual brains and are distributed in the bilateral sensorimotor cortex, with a clear lateralization towards the right (ipsilateral) hemisphere. However, the training effect has a more robust and stable neural representation centered in the central sulcus, while the transfer effect shows weaker but reliable neural codes, including in the central sulcus, but mainly in the surrounding sensorimotor territories. Moreover, the motor engrams shaped by these different learning processes have distinct representations in the sensorimotor cortex, also shifted towards the right hemisphere, particularly in the right side of the superior part of the precentral sulcus, postcentral gyrus, postcentral sulcus, and in the left inferior part of the precentral sulcus. Altogether, our findings show that practice-based learning and transfer/generalization-based learning have distinct representations embedded in spatially overlapping territories, which lead to diverging motor engrams. Finally, these neural codes of motor learning follow a shared high-dimensional structure across individual brains.

## Results

We used 7-Tesla fMRI data to assess the multivariate patterns of brain responses to sequential finger-tapping movements of trained and untrained sequences. The participants (n=13, 9 females, mean age of 21.4 (std=2.36)) performed finger-tapping movements of 12 different sequences before and after a training protocol of five weeks, during which six of them were practiced for around 4000 trials (9). All sequences were matched by their difficulty level and consisted of nine-digit-long sequences that were performed at the same pace with the five fingers of the right hand. Each sequence was executed twice in a row, three times within a run. Each session consisted of eight runs.

We used hyperalignment to capture the response basis functions that are shared across individuals and project them into a common model, which has been shown to lead to above-chance level accuracy in between-subject classification tests (12, 18). To avoid introducing noise to our analysis, we selected the first 12 pairs of trials (six of them of trained sequences, and six of untrained) executed without errors per run (Fig. 1A).

**Figure 1.**
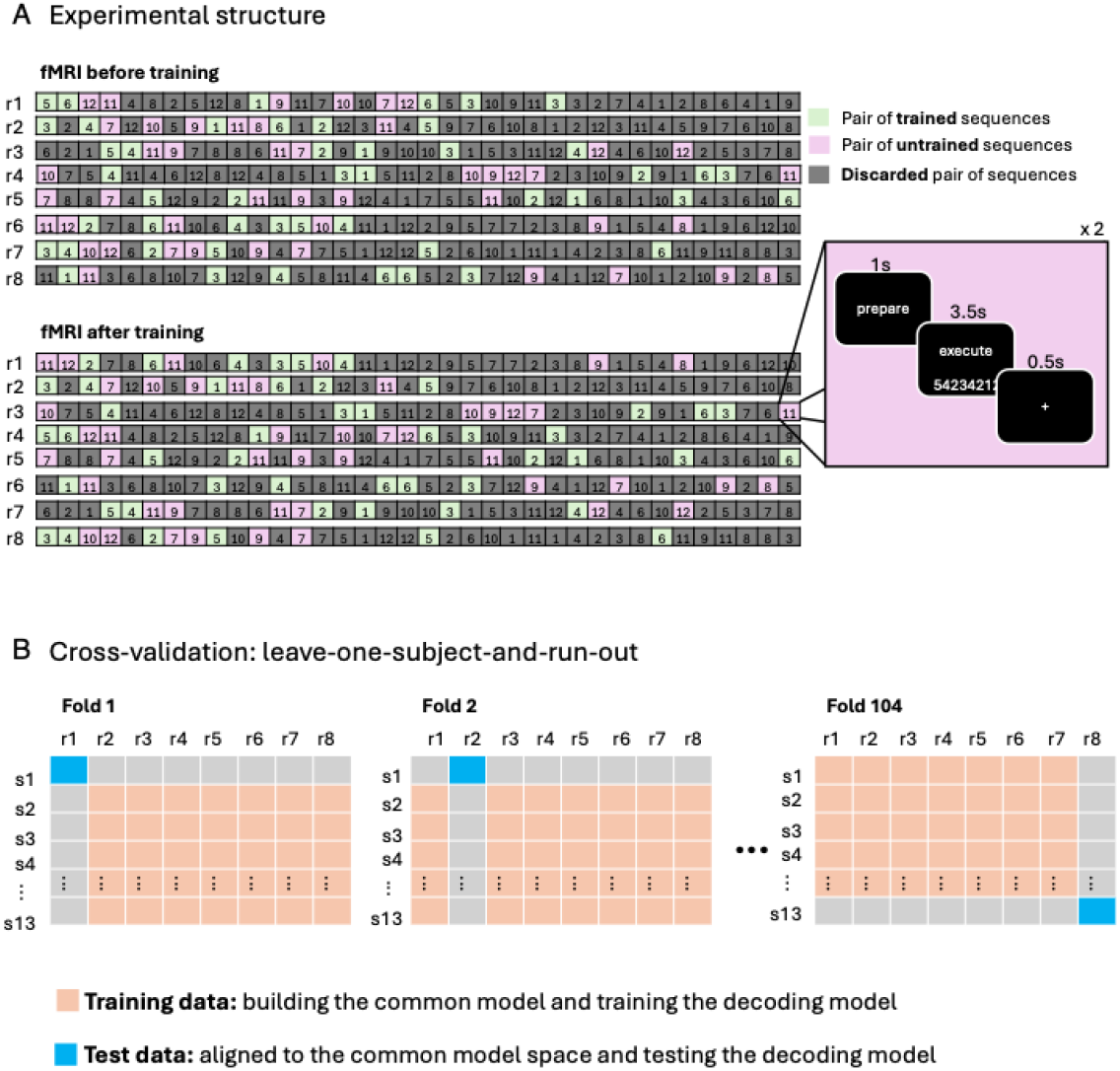
Experimental structure and cross-validation scheme. (A) Before and after the five-week training, participants (s1–s13) were scanned while executing 12 different sequences of finger-tapping movements. Each sequence was performed twice in a row (represented by a single square), three times per run, for eight runs (r1–r8). To reduce noise and task-unrelated signal, the first 12 pairs of sequences (six trained, green; six untrained, purple) were selected and modeled using a surface-based first-level general linear model (GLM), and each square represents the resultant z-map of a pair of trials. Each trial consisted of 1s of preparation, followed by 3.5s of execution (with visual aid) and 0.5s of inter-trial interval. Periods of rest (not shown) were added between trials. (B) The between-subject classification was performed by first setting aside the test subject. The training data (orange) consisted of all runs but one of the remaining subjects, which was used to create the common space and to train the decoder. Then, the left-out run of the test subject (blue) was aligned to the common space and used to test the decoder’s accuracy. We repeated this procedure so that each run from each subject was used as test data.

We built a common model of the sensorimotor cortex (19) using spherical searchlights by first setting aside the data from the test subject (Fig. 1B). The training data consisted of all runs but one of the remaining subjects, which was used to create the common space and train the decoder. Then, the left-out run of the test subject was aligned to the common space and used to test the decoder’s accuracy. We repeated this procedure so that each run from each subject was used as test data (13 subjects x 8 runs = 104 folds). Importantly, we avoided overfitting by keeping apart the training and test data to build the common space and train the decoder (Fig. 1B). This procedure was conducted independently three times, modelling different aspects of motor behavior. In two of them, we investigated the effect of time (before and after motor training) on the neural signatures of the execution of trained and untrained sequences, henceforth called ‘training effect’ and ‘transfer effect’, respectively. In addition, we explored the mechanism-specificity of the neural signatures after the motor training on both trained and untrained sequences, henceforth called ‘resultant motor engrams’, respectively.

A repeated-measures ANOVA revealed that motor training resulted in motor improvements in both trained and untrained sequences, confirming the occurrence of both learning mechanisms. The movement time (Figure 2) of trained sequences decreased from 2.44 s (mean, standard error = 0.13) before the training protocol to 1.3 s (mean, standard error = 0.09) after it, whereas that of the untrained sequences went from 2.46 s (mean, standard error = 0.13) to 1.46 s (mean, standard error = 0.12).

**Figure 2.**
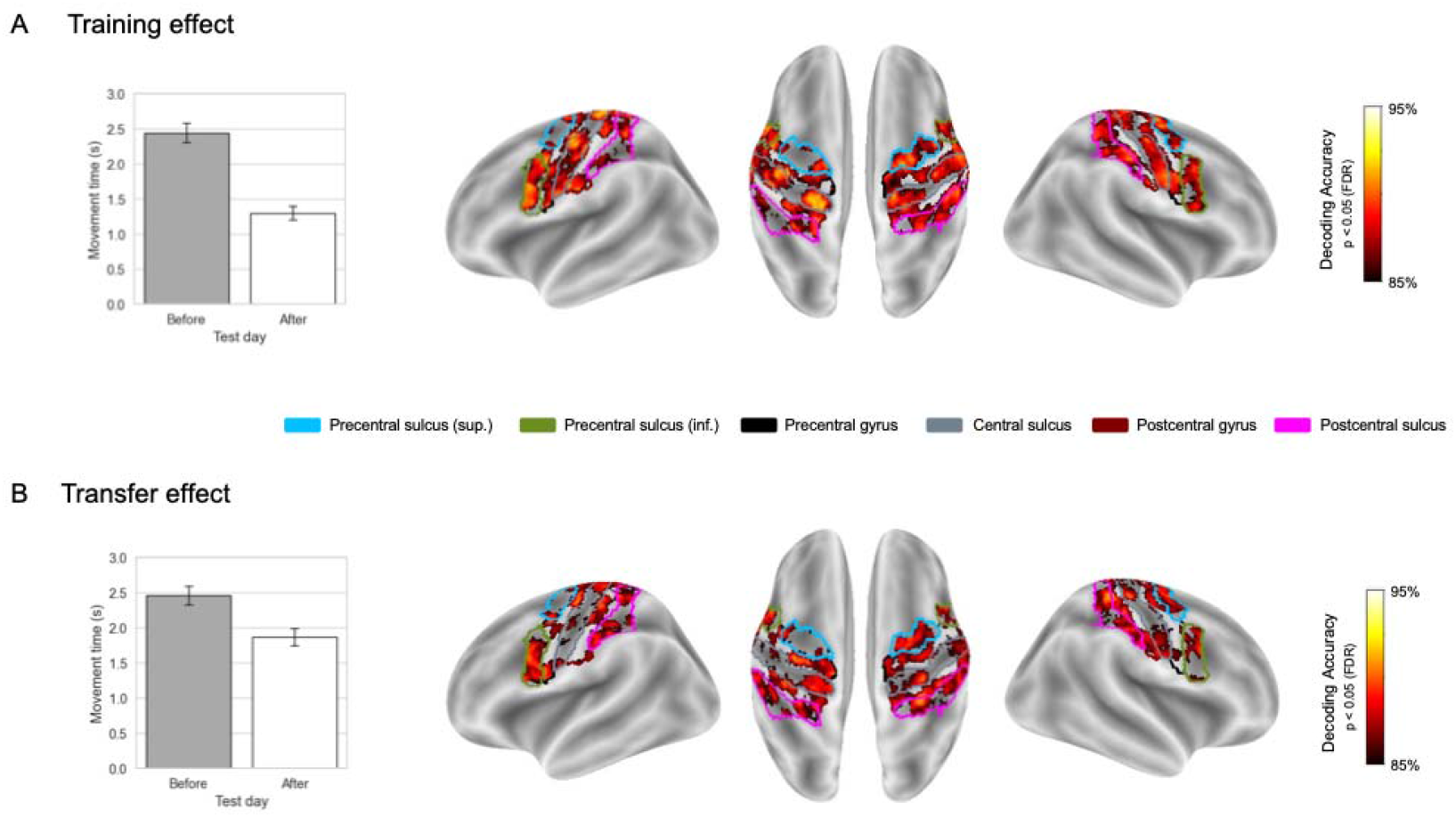
Motor learning and the decoding models for the training and transfer effects. Motor performance and brain maps showing decoding accuracy of the training (A) and transfer (B) effects. Motor learning occurred for both trained (t=10.66; p=1.78e-07) and untrained (t=6.38; p=3.49e-05) sequences. The brain maps show vertices that survived the permutation test (p < 0.05) and FDR correction (Benjamini-Hochberg). Both motor learning mechanisms rely more on the right (ipsilateral) sensorimotor cortex (repeated-measures ANOVA, main effect of side: F(1,12)=8.43,p=0.013). The training effect showed an overall higher accuracy than the transfer effect (main effect of condition: F(1,12)=9.29,p=0.01). In addition, the central sulcus (p=0.0003), the inferior (p = 0.03) and superior (p = 0.03) precentral sulci showed greater decoding accuracy in the training effect than the transfer effect (interaction of condition*label: F(5,60)=15.68, p<0.000). Both left and right central sulci (p = 0.0002 and 0.008, respectively) and the inferior part of the right precentral sulcus showed higher decoding accuracy in the training than the transfer effect (interaction condition*label*side: (F(5,60)=3.88,p=0.004)). See SI.

### The training and transfer effects are embedded in similar territories of the sensorimotor cortex, but with distinct neural codes

The investigation of the neural codes of the training and transfer revealed a bilateral engagement of multiple areas (Figure 2; Table 1) within the sensorimotor cortex. We compared the average decoding accuracy of suprathreshold vertices in each of the six cortical labels in each hemisphere (repeated-measures ANOVA, within-subject factors: condition, cortical label, and side; See SI). The training effect showed an overall higher accuracy than the transfer effect, especially in the bilateral central sulcus and the inferior part of the right precentral gyrus. Interestingly, the central sulcus showed a clear divergent involvement in both mechanisms (Figure S1), consistent with the hierarchical organization of motor learning in the sensorimotor cortex (8). This key distinction was also observed by contrasting both accuracy maps (Training vs. Transfer; Figure 3A), as the bilateral central sulci showed a clear mechanism-specificity, in contrast to the rest of the sensorimotor cortex.

**Table 1.** Average decoding accuracy (training and transfer effects)

| Label | Training effect |  | Transfer effect |  |
| --- | --- | --- | --- | --- |
|  | Left | Right | Left | Right |
| Central sulcus** | 88.029 (0.276)* | 87.609 (0.262)* | 85.974 (0.285) | 86.001 (0.286) |
| Postcentral gyrus | 86.522 (0.28) | 86.381 (0.261) | 86.257 (0.278) | 86.988 (0.28) |
| Postcentral sulcus | 86.043 (0.307) | 87.447 (0.261) | 86.571 (0.314) | 87.07 (0.296) |
| Precentral gyrus | 87.161 (0.278) | 86.776 (0.271) | 87.026 (0.268) | 86.155 (0.267) |
| Precentral sulcus (inf.)** | 87.897 (0.284) | 87.031 (0.294)* | 87.286 (0.293) | 86.064 (0.343) |
| Precentral sulcus (sup.)** | 85.811 (0.324) | 88.411 (0.279) | 85.169 (0.308) | 87.5 (0.264) |

Asterisks highlight statistically significant (FDR-corrected $p < 0.05$ ) interaction of condition x side x label (\*) and condition x label (\*\*).

**Figure 3.**
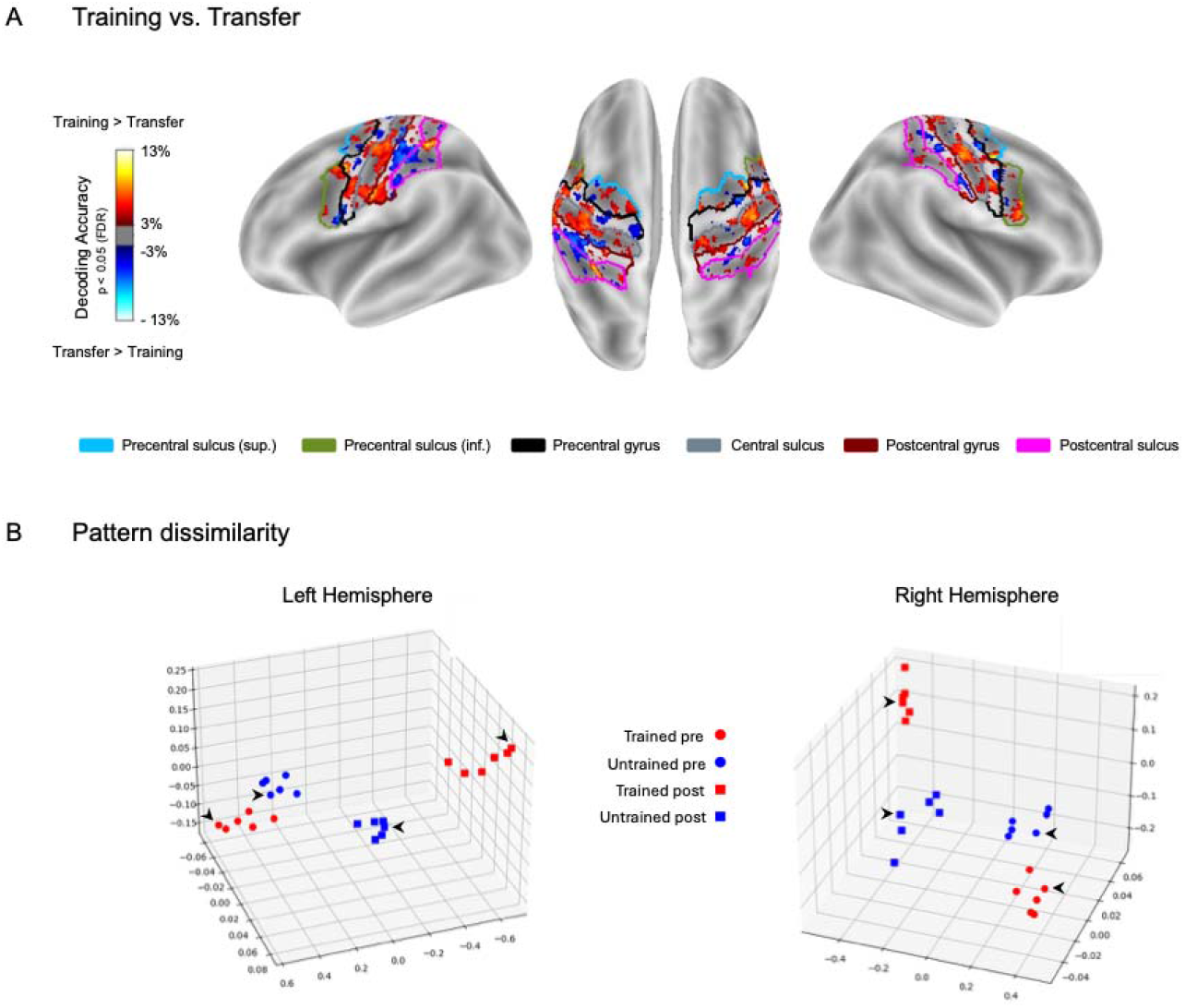
Direct comparison of the neural codes of training and transfer effects with hyperalignment. (A) Red to yellow vertices indicate higher decoding accuracy for the training effect (Training > Transfer), while blue to cyan vertices indicate higher decoding accuracy for the transfer effect (Transfer > Training). The map shows vertices that survived a two-sided test, permutation testing, and FDR correction (Benjamini-Hochberg). (B) The multidimensional scaling plot displays the dissimilarity in all sensorimotor cortical labels, showing that after motor training (squares), the representational patterns of both trained (red) and untrained sequences (blue) diverge from their pre-training configuration (blue and red circles), with trained sequences exhibiting a more pronounced shift. Arrowheads highlight the central sulcus.

A multidimensional scaling plot (Figure 3B) illustrates that, after motor training, the representational patterns of both trained and untrained sequences diverge from their pre-training configuration, with trained sequences exhibiting a more pronounced shift.

### The resultant motor engrams are mechanism specific

After the motor training protocol, the mechanism by which motor learning occurred was decoded above chance in all cortical labels (Figure 4, Figure S1; Table 2), with a clear overall shift towards the right hemisphere, and specifically in the superior part of the precentral sulcus, the postcentral sulcus and gyrus. In contrast, the inferior part of the precentral sulcus showed higher decoding accuracy in the left hemisphere. Within the left hemisphere, the inferior part of the precentral sulcus showed the highest decoding accuracy. In the right hemisphere, the superior part of the precentral sulcus showed higher decoding accuracy than the precentral and postcentral gyri, and the postcentral sulcus showed higher decoding accuracy than the postcentral gyrus.

**Table 2.** Average decoding accuracy (resultant motor engrams)

| Label | Left | Right |
| --- | --- | --- |
| Central sulcus | 74.435 (0.29) | 74.386 (0.305) |
| Postcentral gyrus* | 73.977 (0.28) | 74.38 (0.285) |
| Postcentral sulcus* | 73.776 (0.29) | 74.753 (0.3) |
| Precentral gyrus | 74.15 (0.284) | 74.391 (0.305) |
| Precentral sulcus (inf.)* | 75.138 (0.317) | 74.701 (0.311) |
| Precentral sulcus (sup.)* | 73.842 (0.284) | 75.065 (0.315) |

\* Statistically significant (FDR-corrected $p < 0.05$ ) label x side interaction.

**Figure 4.**
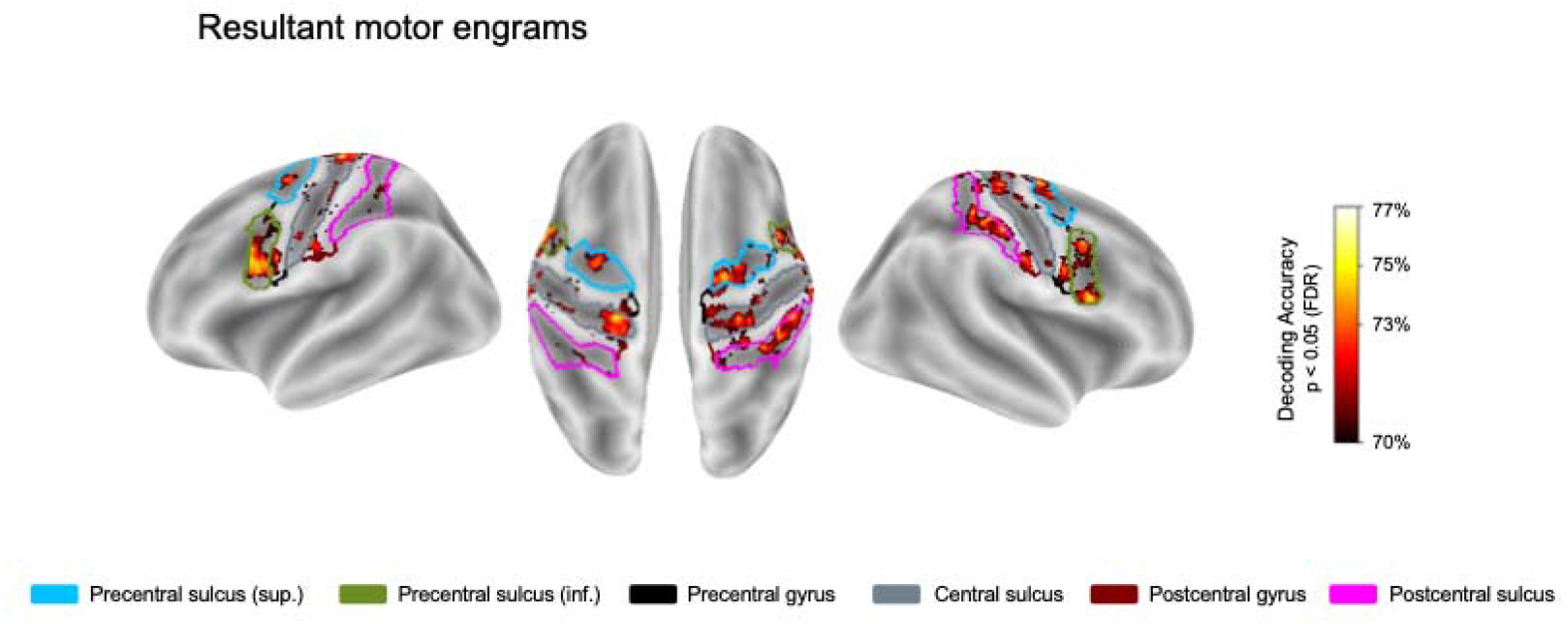
Decoding model for the resultant motor engrams. Brain maps showing decoding accuracy of the model that compared the resultant motor engrams, revealing that the resultant motor engrams were decoded above chance across all labels, with stronger encoding in the right hemisphere. The brain map shows vertices that survived the permutation test (p < 0.05) and FDR correction (Benjamini-Hochberg). A repeated-measures ANOVA showed that the decoding accuracy is greater in the right hemisphere (main effect of side: F(1,12)=21.25,p=0.0006), and specifically (interaction label*side) in the superior part of the precentral sulcus (t=-10.11, p=0.000), the postcentral sulcus (t=- 5.42, p=0.000) and gyrus (t=-2.83, p=0.03); and in the inferior part of the precentral sulcus (t=2.46, p=0.04) of the left hemisphere. Within the left hemisphere, the inferior part of the precentral sulcus showed the highest decoding accuracy. In the right hemisphere, the superior part of the precentral sulcus showed higher decoding accuracy than the precentral (t=-3.87, p=0.03) and postcentral (t=-3.61, p=0.03) gyri, and the postcentral sulcus showed higher decoding accuracy than the postcentral gyrus (t=-3.37, p=0.03).

## Discussion

In the present study, we investigated the high-dimensional representation of the training-based and transfer/generalization-based learning of sequential finger-tapping movements. We found that (i) both learning processes have distinct neural representations, even though embedded in a spatially overlapping cortical landscape, which (ii) lead to two distinct motor engrams. Importantly, both motor learning mechanisms and their resultant motor engrams are encoded following the same high-dimensional architecture across individual brains. The neural representations of the training and transfer effects are distributed bilaterally, but more pronouncedly in the right (ipsilateral) sensorimotor cortex. The training effect showed an overall higher decoding accuracy than the transfer effect, suggestive of a robust neural representation. Interestingly, the central sulcus displayed distinct encoding processes, clearly marking its differential roles in motor learning by repetitive training and generalization. Finally, the neural processes underlying the training and transfer effects lead to divergent motor engrams. Altogether, our findings support previous studies showing that both practice-based and transfer/generalization-based motor learning recruit similar cortical structures (2–5, 20). Despite the spatial similarity, we extend previous findings by demonstrating that these two learning processes encode different neural content, resulting in divergent motor engrams, which emphasizes the idea that training and transfer involve separate mechanisms. Reinforcing the model of hierarchical organization of motor skill learning, these findings suggest that repetitive training consolidates rigid, execution-focused representations in primary sensorimotor areas, whereas transfer or generalization relies mostly on higher-level representations outside them.

Studies have shown that the sensorimotor cortex is differentially engaged over the course of a motor training protocol and in response to the transfer of learning, showing either activation increases and decreases, depending on the stage of the learning process (1, 9, 20–22). In motor learning, increased brain activity is commonly interpreted as the strengthening of neural circuits involved, whereas decreased activity is often taken as neural optimization or better processing efficiency (22–24). However, these fluctuations are difficult to interpret, as they likely reflect the summation of multiple complex neural processes occurring during motor learning (25). Moreover, simply reporting that certain brain regions display higher or lower activation after training does not inform about the specific nature of their contributions to the task – it merely describes the extent of involvement, not its content. Therefore, investigating the content of the cortical engagement during practice-based learning and transfer/generalization-based learning requires multivariate methods capable of decoding ‘*what’* is encoded (26). In this way, if the neural circuitry of the training and transfer effects were identical (if both learning mechanisms were, in fact, a single phenomenon), it would not be possible to distinguish them at the neural level. In contrast to that, we deepen the comprehension of the neural basis of motor learning by describing that the sensorimotor cortex is differentially involved in those processes.

The anatomical variability and idiosyncratic functional response of cortical structures are the main challenges that hamper between-subject classification (27) of neurophysiological data. Functional alignment using hyperalignment overcomes this by creating a high-dimensional model that represents the response-tuning functions that are common across individual brains (12, 26). In this type of approach, the cross-validation scheme is of extreme importance. Here, we avoided data leakage by keeping the train and test data apart for the common model estimation and decoder training. Thus, we can conclude that the only possible way to get above-chance decoding accuracy in our between-subject classification model is if both training and test data contain similar representation structures. Hence, our findings revealed the existence of a shared, high-dimensional information space that represents motor learning achieved by two distinct mechanisms. This is in line with previous studies showing that the sensorimotor representation of finger-tapping movements (18) follows a similar structure across individual brains. Interestingly, such a procedure was not effective using standard, anatomical alignment (Fig. S4, see SI).

We showed that the training effect relies robustly on the central sulcus, diverging from what happens in the transfer effect. This is consistent with the view that motor skill learning is structured in layers, from high-level action plans to low-level muscle commands (8). Here, repetitive practice likely sculpted execution-level primitives in the primary sensorimotor cortex, reflecting the formation of stable, invariant neural codes that automate execution, which lead to higher decoding accuracy across M1/S1. In contrast, transfer/generalization encodes more abstract sequence representations outside the central sulcus. This interpretation is supported by our direct comparison between both learning mechanisms, which might indicate the formation of specialized and more stable neural circuits through repetitive training in the sensorimotor cortex, especially in M1/S1 (8, 9, 28, 29), whereas the transfer effect showed weaker but still reliable decoding accuracy in more rostral and causal areas, representing the variability and less informative characteristics of higher-level sequence representations (9, 11, 28–30).

Motor engrams are known to support the execution and acquisition of skilled movements (31). Here, we showed that the motor engrams sculpted by repetitive training and generalization are distinct. These engrams were sufficiently distinct to be decoded with above-chance accuracy in all the sensorimotor cortical regions. Interestingly, the right (ipsilateral) hemisphere showed greater contribution to the decoding models of each learning process and their resultant motor engrams than the left hemisphere. While ipsilateral activity of the sensorimotor cortex has been described in multiple studies (2, 3, 10, 28, 32–36), its role in motor learning is still under debate. According to the “interhemispheric competition” model, in which the ipsilateral sensorimotor cortex receives inhibitory inputs from the contralateral hemisphere (37), the ipsilateral brain activity during unimanual motor tasks is associated with impaired motor behavior, and suppressing it is beneficial for motor learning in healthy individuals (38, 39) and motor recovery after stroke (40–42). Likewise, stimulating ipsilateral M1 impairs motor learning (43). In contrast to this view, other studies point to a collaborative nature of ipsilateral activity that may occur through changes in the interhemispheric communication (44). Even though weaker, the ipsilateral sensorimotor cortex encodes finger movements (45, 46) and finger-tapping sequences (47), and stimulating these areas has been linked to a positive impact on motor learning (48).

In conclusion, our study shows that the training and transfer effects on motor skill learning have different neural codes embedded in a similar cortical topography, which are shared across individual brains. The training and transfer effects shape distinct motor engrams, reinforcing the idea that they represent different motor learning processes. Importantly, our results highlighted the key engagement of the ipsilateral sensorimotor cortex in motor learning encoding, which deserves further investigation in future studies.

## Materials and Methods

The present study is a secondary analysis of a publicly available dataset (OpenNeuro, accession number ds002776) that has been published elsewhere (9). The experimental description below was obtained from publicly available resources, such as the original publication (9) and the dataset description. Specific parts irrelevant to the present study (e.g., behavioral test sessions outside the MRI, free-speed execution) have been omitted for clarity purposes. For a full description of the experimental design, refer to the original publication (9).

### Participants

In the present study, we used fMRI and behavioral data of 13 healthy participants (9 females, mean age of 21.4 (sd=2.36)) who were part of group 1 in the original dataset, and after exclusion of one participant (sub-24) due to the high number of errors during the task-based fMRI. All participants were right-handed, had no prior history of psychiatric or neurologic disorders, and provided written informed consent to all procedures and data usage. The experimental procedures were approved by the Ethics Committee at Western University, where the data has been collected.

### Training protocol

The training protocol consisted of training the execution of six 9-digit finger sequences, for five weeks, as fast as possible. Participants received feedback based on the accuracy and movement time, both online and after each trial, respectively. Each sequence was executed twice in a row, first with visual cues indicating the finger-tapping sequence, then without visual aid. Each training block consisted of 24 trials (4 trials per trained sequence) and was alternated with rest periods. The training protocol occurred 3 days per week in the first 4 weeks, and on one day in the last week. The trained sequences have been practiced for around 4000 trials in total.

### Imaging acquisition

MRI data have been acquired using a 7-Tesla Siemens Magnetom scanner with a 32-receive channel head coil (8-channel parallel transmit). Anatomical T1-weighted scan used a magnetization-prepared rapid gradient echo sequence (MPRAGE) with a voxel size of 0.75 x 0.75 x 0.75 mm isotropic (field of view = 208 x 157 x 110 mm [A-P; R-L; F-H], encoding direction coronal). For fMRI data, the GRAPPA3 sequence (multiband acceleration factor 2, repetition time [TR]=1.0 s, echo time [TE]=20 ms, flip angle [FA]=30 deg, 44 slices with isotropic voxel size of 2 x 2 x 2 mm) has been used. Gradient echo field map has been collected to estimate magnetic field inhomogeneities (transversal orientation, field of view 210 210 160 mm and 64 slices with 2.5 mm thickness, TR = 475 ms, TE = 4.08 ms, FA = 35 deg).

### Experimental design

In the present study, we analyzed data from two different scanning sessions: one before and one after the five weeks of training. Each scanning session consisted of eight functional runs with an event-related design, randomly intermixing execution of trained and untrained sequences. Each sequence was repeated twice in a row, for three times within a run. Trials started with 1 s preparation time, during which the sequence was presented on the screen. After that time, a ‘go’ signal was displayed as a short pink and expanding line underneath the sequence numbers that indicated the speed at which participants were required to press along. The execution phase, including the feedback on overall performance, lasted for 3.5 s, and the inter-trial interval was 0.5 s. Each trial lasted for 5 s. Five periods of rest, each 10 s long, were added randomly between trials in each run.

### Data preprocessing

Data has been preprocessed using fmriprep version 24.0.0. T1w images were corrected for intensity non-uniformity and used as T1w-reference throughout the workflow. The T1w-reference was then skull-stripped, and brain tissue segmentation of cerebrospinal fluid (CSF), white-matter (WM), and gray matter (GM) was performed. Non-linear spatial normalization to the standard space (MNI152NLin2009cAsym) was performed using brain-extracted versions of both T1w reference and the T1w template. Head motion parameters, including six rotation and translation components and transformation matrices, were estimated relative to this reference image before any spatiotemporal filtering.

Subsequently, the BOLD reference image was co-registered to the T1-weighted (T1w) anatomical reference using boundary-based registration with six degrees of freedom. Framewise displacement (FD) and DVARS metrics were calculated for each functional run. Additionally, mean signals were extracted from cerebrospinal fluid (CSF), white matter (WM), and whole-brain masks. Temporal high-pass filtering (using a discrete cosine filter with a 128-second cutoff) was applied to the preprocessed BOLD time series before principal component analyses for both anatomical (aCompCor) and temporal (tCompCor) CompCor decompositions. For aCompCor, probabilistic tissue masks for CSF, WM, and a combined CSF+WM mask were generated in anatomical space. Additional nuisance regressors included temporal derivatives and quadratic expansions of motion parameters and global signals.

## Data analysis

After preprocessing, BOLD images were analyzed using Python libraries, such as nilearn, neuroboros, nibabel, and nltools. The surface-based first-level general linear model (GLM) was implemented using Nilearn’s FirstLevelModel and incorporated task-related regressors for each condition, confound regressors, and an SPM hemodynamic response function (HRF) model. An autoregressive (AR1) noise model was specified. The confound regressors included WM signal, global signal, FD, and six motion parameters (translational and rotational), which were calculated with fmriprep. Task timing and event details were extracted from event files, which specified the onset, duration, and condition of each trial. Each trial (modeled with a 3.5s length) was assigned a unique condition label combining sequence type (trained or untrained), sequence number (1-12), and pair of repetitions (from 1 to 3). In each run, contrasts were computed to estimate task-related activation. The resulting z-maps for each condition (e.g., trained_6_pair1) were then masked for the sensorimotor cortex (19) and used for the decoding analysis. In order to consider only perfectly executed trials but keeping the homogenous and balanced data structure, we selected the first 12 conditions (6 trained, 6 untrained) of each participant and run. The remaining data was discarded and not used in our analyses.

### Searchlight hyperalignment

Inside each disk of radius 20 mm, hyperalignment was performed to align the neural representations across participants to a common representational space. For each fold in the leave-one-subject-and-run-out cross-validation scheme, all runs but one from the training data were used to construct the representation space. This was achieved using a Procrustes-based algorithm, which iteratively transforms the training data into a shared representational space (common space) while preserving the geometry of individual neural patterns (27). After creating the hyperalignment model inside each cortical disk using the training data, the test data (the left-out run of the left-out subject) was projected into the common space. This ensured that the test data were represented within the same common space as the training data, enabling direct comparisons. We repeated this scheme independently for each decoding model (training effect, transfer effect, resultant motor engrams) in such a way that each run of each subject was used as test data once, resulting in 104 folds.

### Searchlight decoder implementation

A searchlight-based linear support vector machine (SVM) with L1 regularization was used as the decoding algorithm to classify neural activity patterns in cortical disks of radius 10 mm. The decoder was trained on the hyperaligned training data and their corresponding labels, and then tested on the hyperaligned test data. We used principal component analysis for dimensionality reduction inside each searchlight. Classification accuracies from each searchlight were placed into their center surface nodes, resulting in one accuracy map for each cross-validation fold. We used permutation testing for statistical thresholding by first repeating the analysis 100 times with randomly shuffled sequence identity labels in each searchlight. For each hemisphere and vertex, we aggregated individual-level decoding accuracies from cross-validation folds. The empirical null distribution was obtained by sampling decoding accuracies from permutation-based analyses (i.e., models tested on data with randomized labels). For each vertex, we drew bootstrap samples randomly with replacements from the pool of permutation scores. We repeated this procedure 10,000 times, each time computing the mean of the sampled values to create a distribution of group-level means under the null hypothesis. Observed decoding performance at each vertex was then compared against its corresponding null distribution. The empirical P values were computed by counting the proportion of null distribution values greater than or equal to the observed decoding accuracy, adding one to both the numerator and denominator to avoid zero probabilities and reduce bias. This constitutes a one-sided non-parametric test for above-chance performance. The Benjamini–Hochberg procedure was subsequently applied to correct P values for multiple comparisons.

A similar approach was applied to compute the difference in classification accuracy between the training and transfer effects: we averaged the accuracy values from the 104 cross-validation folds for the two models independently, and then set accuracy values below the chance level to 50% to avoid noise and artificial inflation of the difference (17). The same procedure was applied 10,000 times to the permuted cross-validation folds, generating a null distribution of difference values for each searchlight. A two-sided P value was computed by counting how many times the absolute value of the null distribution exceeded the absolute value of the original unpermuted difference map (adding 1 to both the numerator and denominator). The Benjamini– Hochberg procedure was subsequently applied to correct P values for multiple comparisons

## Statistical Analysis

For each participant, the mean accuracy per cortical area was calculated and repeated-measures ANOVAs were performed on the mean accuracy (dependent variable) with condition (trained, untrained), cortical label (anterior and superior parts of the precentral sulcus, precentral gyrus, central sulcus, postcentral sulcus, and postcentral gyrus) and side (left and right hemispheres) as within-subjects factors, and subjects as the random factor.

Using hyperalignment, the decoding models led to significant findings in most vertices after permutation testing and FDR (Benjamini–Hochberg) correction. The accuracy values closest to zero that passed the statistical threshold were: 75.4% and 69.9% for the training effect; 73% and 75.1% for the transfer effect; 0.12% (positive) /-0.12% (negative) and 0.04% (positive) / - 0.08% (negative) for the difference between them, and 60.3% and 59.1% for the comparison of the resultant motor engrams, in the right and left hemispheres, respectively. As mentioned, the mean accuracy was estimated on the statistically significant vertices only, and the accuracy maps were thresholded arbitrarily to improve plot visualization.

A repeated-measures ANOVA was conducted to compare the average decoding accuracy per cortical label. Post hoc tests were performed using FDR correction, when appropriate.

Pattern dissimilarity among the four condition/session patterns was quantified using the correlation distance, computed as 1 − Pearson correlation, producing a 4x4 dissimilarity matrix per ROI and hemisphere. The dissimilarities were embedded into a three-dimensional configuration using classical metric multidimensional scaling (MDS), preserving the precomputed correlation distances across the four points to visualize relative pattern geometry across trained/untrained and pre/post states.

## Data Availability

The fMRI data is publicly available (OpenNeuro, accession number ds002776). The codes used in this publication will be made publicly available after peer review and publication.

## Acknowledgements

This work was supported by the Austrian Science Fund FWF (ESP 397-BBL).

## Supplementary Information

### Supplementary Results

**Figure S1.**
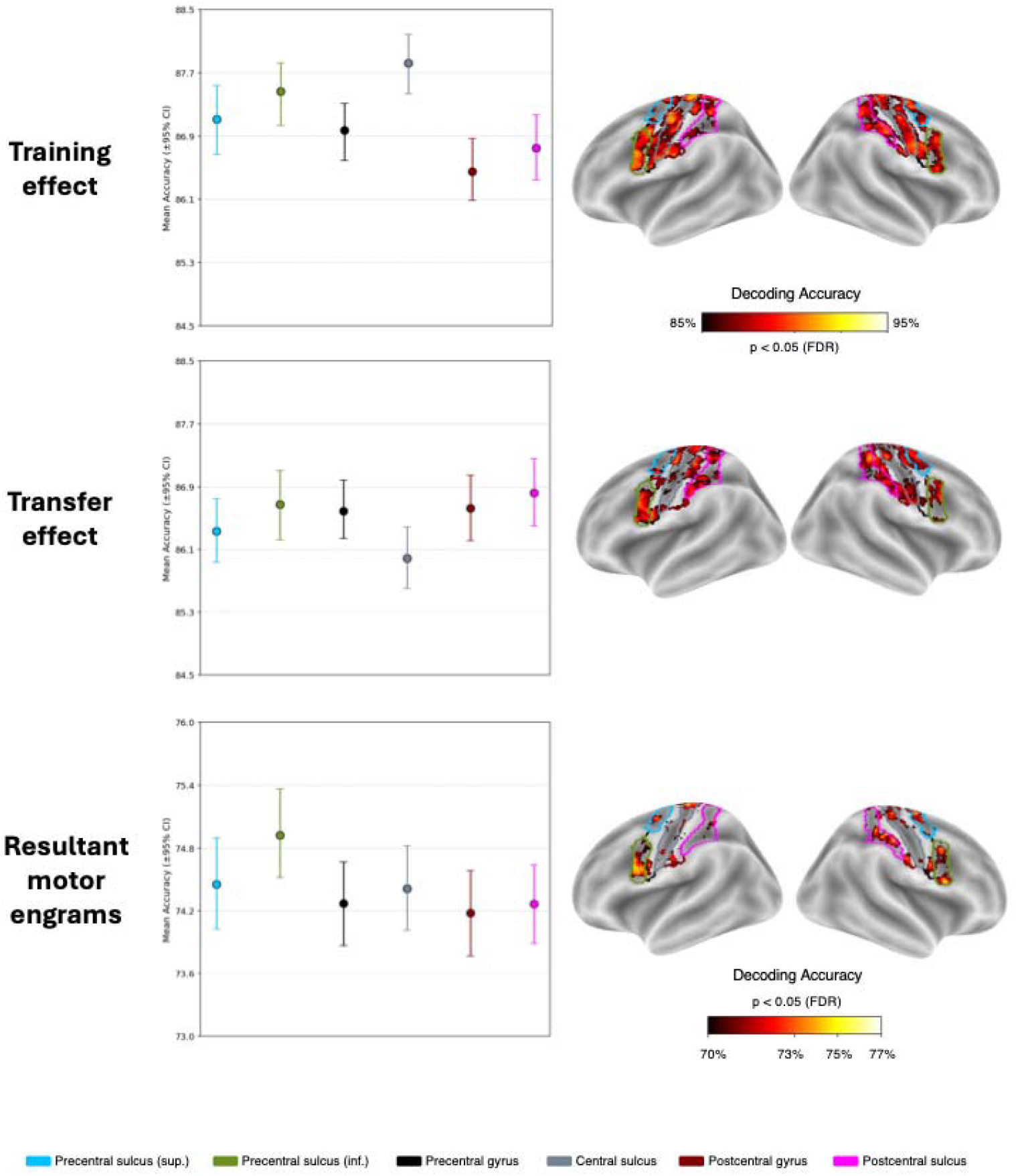
Mean accuracy and confidence interval (95%) obtained with bootstrapping. For each participant, we calculated the mean accuracy per cortical area and estimated the group-level 95% confidence interval based on bootstrapping (random sampling, k = 1,000 with replacement).

**Figure S2.**
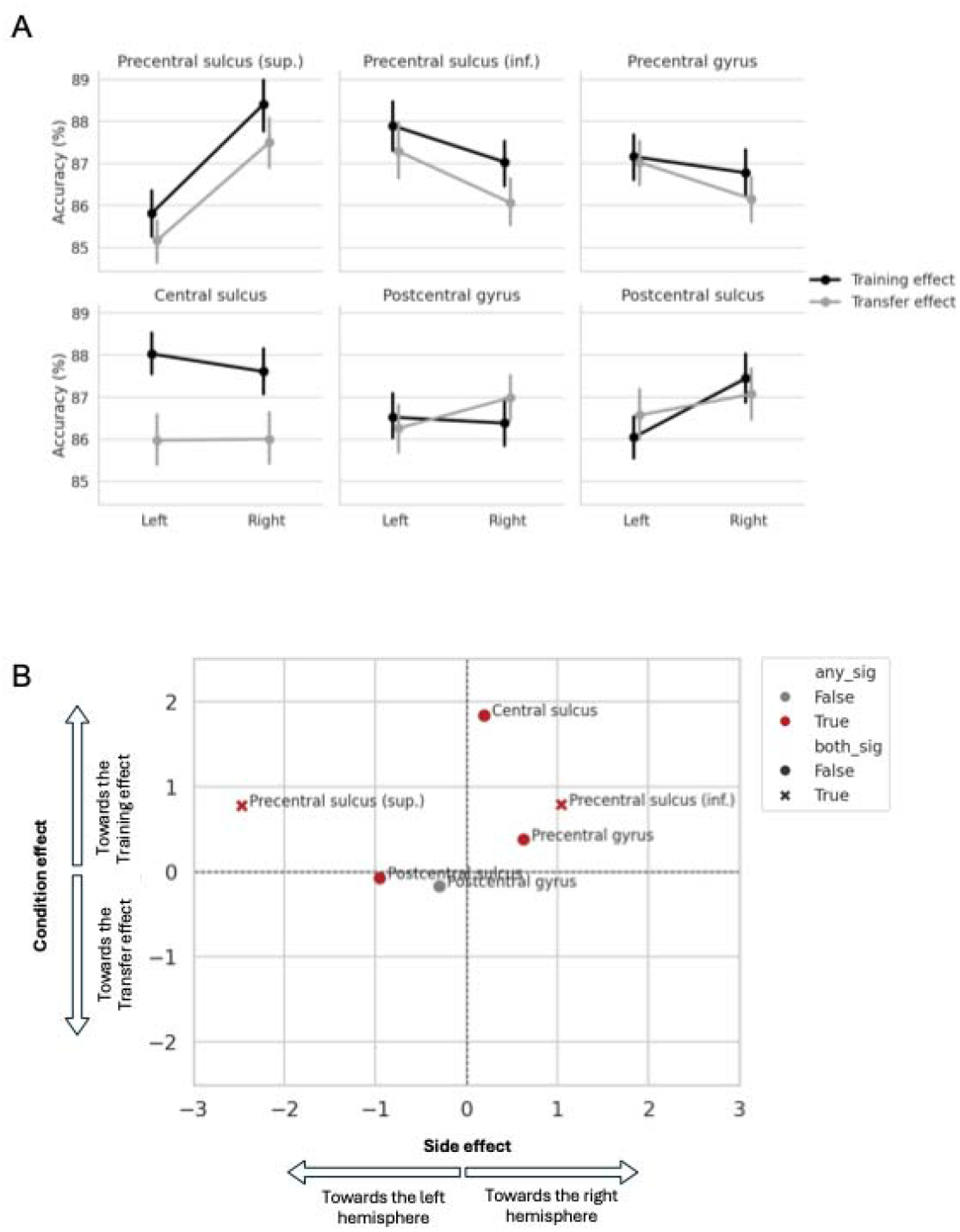
Average decoding accuracy per label. (A) Mean accuracy by side for each anatomical label, with hue indicating condition (Training effect vs Transfer effect) and vertical error bars displaying 95% bootstrap confidence intervals. (B) Two-dimensional effects map showing per-label paired differences in accuracy for Side (left and right hemispheres) and Condition (training and transfer effects) contrasts. Each point represents the mean within-subject difference after collapsing over the other factor, and the vertical and horizontal dashed lines mark zero effect. Points to the left indicate Left > Right, and points above zero indicate Training > Transfer. Color denotes labels significant (p < 0.05) after FDR correction for either contrast, and marker shape denotes labels significant for both contrasts.

**Figure S3.**
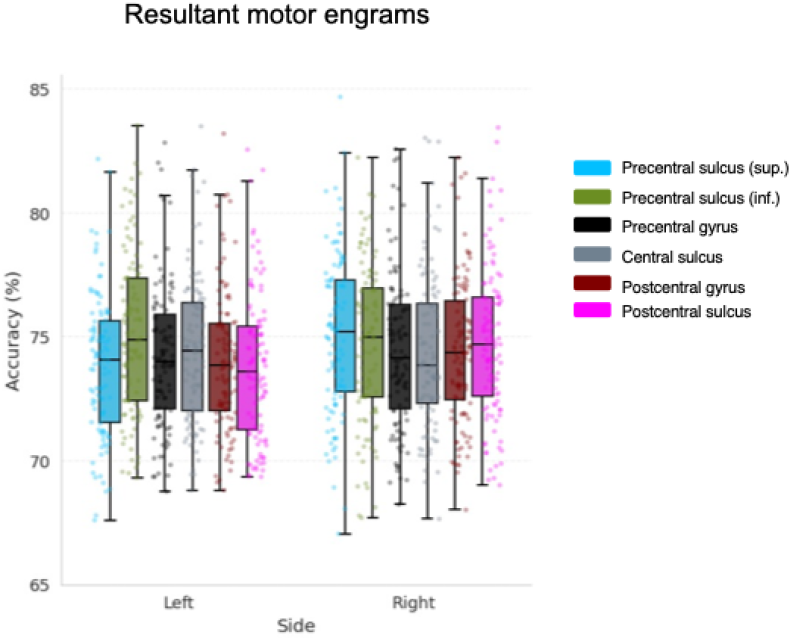
Average decoding accuracy by label and hemisphere of the decoding model that compared the resultant motor engrams. Boxes show group distributions per Side and label (quartiles with whiskers), while overlaid jittered points display individual observations

**Figure S4.**
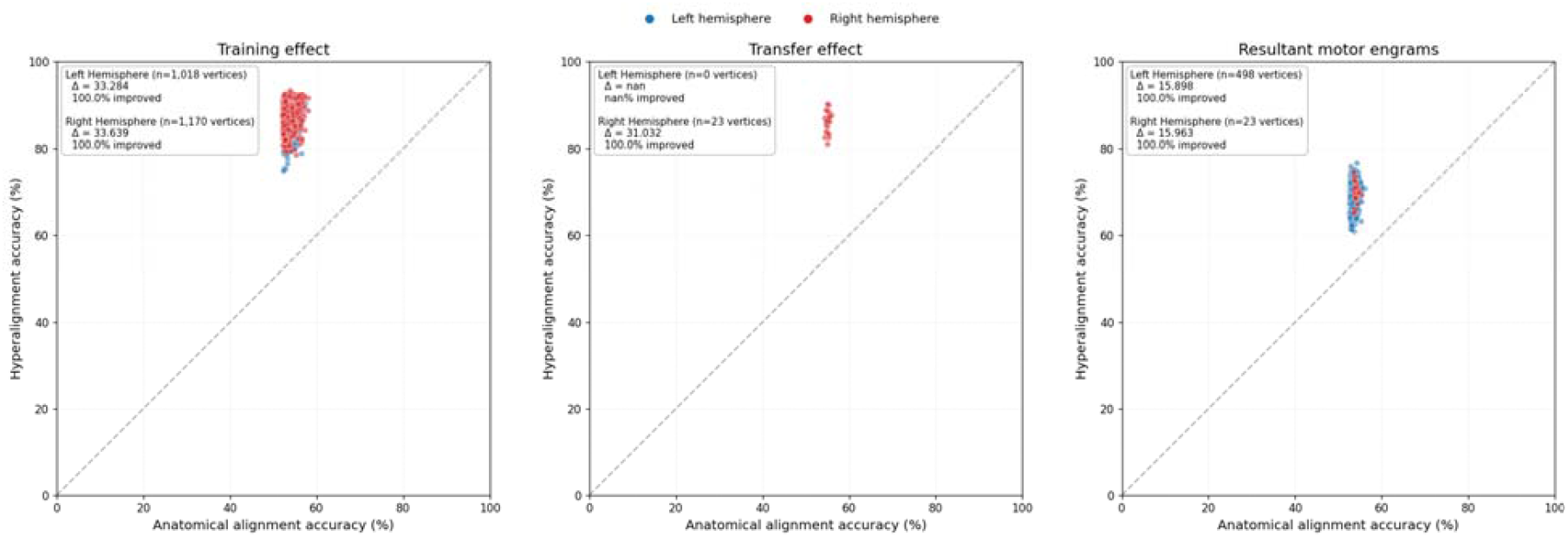
Scatterplot of searchlights’ accuracies for each decoding model using either hyperalignment or standard, anatomical alignment. The between-subject decoding models with anatomically aligned data followed the same procedure as their counterparts on hyperaligned data, except that the individual functional data were kept in the template surface fsaverage. When compared to the decoding models that used hyperaligned data, between-subject decoding models with anatomically aligned data revealed limited decoding accuracy. Within the left (9519 vertices) and right (9506 vertices) hemispheres, the anatomically aligned models returned 1,023 (left) and 1,168 (right) supra-threshold vertices for the training effect, 0 (left) and 23 (right) for the transfer effect, and 498 (left) and 23 (right) for the resultant motor engrams. Across all models, hyperalignment improved decoding accuracy in 100% of the vertices.

## Notes

https://openneuro.org/datasets/ds002776

### Competing Interest Statement

The authors have declared no competing interest.

### Summary of Updates

An overall text revision has been performed with the inclusion of a new coauthor (FT-M). No substantial change in the scientific content has occurred.

## References

1. J. W. Krakauer, A. M. Hadjiosif, J. Xu, A. L. Wong, A. M. Haith, Motor learning. Compr. Physiol. 9, 613–663 (2019).

2. S. Pau, G. Jahn, K. Sakreida, M. Domin, M. Lotze, Encoding and recall of finger sequences in experienced pianists compared with musically naïve controls: A combined behavioral and functional imaging study. NeuroImage 64, 379–387 (2013).

3. R. E. Shimizu, A. D. Wu, B. J. Knowlton, Cerebellar activation during motor sequence learning is associated with subsequent transfer to new sequences. Behav. Neurosci. 130, 572–584 (2016).

4. R. D. Seidler, D. C. Noll, Neuroanatomical Correlates of Motor Acquisition and Motor Transfer. J. Neurophysiol. 99, 1836–1845 (2008).

5. J. Langan, R. D. Seidler, Age differences in spatial working memory contributions to visuomotor adaptation and transfer. Behav. Brain Res. 225, 160–168 (2011).

6. G. Torres-Oviedo, A. J. Bastian, Seeing Is Believing: Effects of Visual Contextual Cues on Learning and Transfer of Locomotor Adaptation. J. Neurosci. 30, 17015–17022 (2010).

7. G. Torres-Oviedo, A. J. Bastian, Natural error patterns enable transfer of motor learning to novel contexts. J. Neurophysiol. 107, 346–356 (2012).

8. J. Diedrichsen, K. Kornysheva, Motor skill learning between selection and execution. Trends Cogn. Sci. 19, 227–233 (2015).

9. E. Berlot, N. J. Popp, J. Diedrichsen, A critical re-evaluation of fMRI signatures of motor sequence learning. eLife 9, 1–24 (2020).

10. A. Yokoi, S. A. Arbuckle, J. Diedrichsen, The Role of Human Primary Motor Cortex in the Production of Skilled Finger Sequences. J. Neurosci. 38, 1430–1442 (2018).

11. A. Yokoi, J. Diedrichsen, Neural Organization of Hierarchical Motor Sequence Representations in the Human Neocortex. Neuron 103, 1178-1190.e7 (2019).

12. J. V. Haxby, J. S. Guntupalli, S. A. Nastase, M. Feilong, Hyperalignment: Modeling shared information encoded in idiosyncratic cortical topographies. eLife 9, 1–26 (2020).

13. P. Borovska, B. de Haas, Individual gaze shapes diverging neural representations. Proc. Natl. Acad. Sci. 121, e2405602121 (2024).

14. M. Feilong, S. A. Nastase, J. S. Guntupalli, J. V. Haxby, Reliable individual differences in fine-grained cortical functional architecture. NeuroImage 183, 375–386 (2018).

15. H. Jung, et al., Action features dominate cortical representation during natural vision. [Preprint] (2025). Available at: https://www.biorxiv.org/content/10.1101/2025.01.30.635800v1 [Accessed 4 June 2025].

16. V. Taschereau-Dumouchel, et al., Towards an unconscious neural reinforcement intervention for common fears. Proc. Natl. Acad. Sci. U. S. A. 115, 3470–3475 (2018).

17. M. Visconti di Oleggio Castello, J. V. Haxby, M. I. Gobbini, Shared neural codes for visual and semantic information about familiar faces in a common representational space. Proc. Natl. Acad. Sci. 118, e2110474118 (2021).

18. T. F. Marins, F. A. C. de Azevedo, G. Wood, A common neural architecture for encoding finger movements. [Preprint] (2025). Available at: https://www.biorxiv.org/content/10.1101/2025.09.11.675501v1 [Accessed 12 September 2025].

19. C. Destrieux, B. Fischl, A. Dale, E. Halgren, Automatic parcellation of human cortical gyri and sulci using standard anatomical nomenclature. NeuroImage 53, 1–15 (2010).

20. R. D. Seidler, Neural Correlates of Motor Learning, Transfer of Learning, and Learning to Learn. Exerc. Sport Sci. Rev. 38, 3 (2010).

21. E. Berlot, N. J. Popp, S. T. Grafton, J. Diedrichsen, Combining Repetition Suppression and Pattern Analysis Provides New Insights into the Role of M1 and Parietal Areas in Skilled Sequential Actions. J. Neurosci. 41, 7649–7661 (2021).

22. E. Dayan, L. G. Cohen, Neuroplasticity subserving motor skill learning. Neuron 72, 443–454 (2011).

23. C. Kelly, J. J. Foxe, H. Garavan, Patterns of Normal Human Brain Plasticity After Practice and Their Implications for Neurorehabilitation. Arch. Phys. Med. Rehabil. 87, 20–29 (2006).

24. R. M. Hardwick, C. Rottschy, R. C. Miall, S. B. Eickhoff, A quantitative meta-analysis and review of motor learning in the human brain. NeuroImage 67, 283–297 (2013).

25. C. J. Steele, V. B. Penhune, Specific Increases within Global Decreases: A Functional Magnetic Resonance Imaging Investigation of Five Days of Motor Sequence Learning. J. Neurosci. 30, 8332–8341 (2010).

26. J. V. Haxby, A. C. Connolly, J. S. Guntupalli, Decoding Neural Representational Spaces Using Multivariate Pattern Analysis. Annu. Rev. Neurosci. 37, 435–456 (2014).

27. J. V. Haxby, et al., A Common, High-Dimensional Model of the Representational Space in Human Ventral Temporal Cortex. Neuron 72, 404–416 (2011).

28. T. Wiestler, J. Diedrichsen, Skill learning strengthens cortical representations of motor sequences. eLife 2013 (2013).

29. K. Kornysheva, J. Diedrichsen, Human premotor areas parse sequences into their spatial and temporal features. eLife 3, e03043 (2014).

30. B. Garzón, et al., Cortical changes during the learning of sequences of simultaneous finger presses. Imaging Neurosci. 1, 1–26 (2023).

31. M.-H. Monfils, E. J. Plautz, J. A. Kleim, In Search of the Motor Engram: Motor Map Plasticity as a Mechanism for Encoding Motor Experience. The Neuroscientist 11, 471–483 (2005).

32. I. Nambu, et al., Decoding sequential finger movements from preparatory activity in higher-order motor regions: a functional magnetic resonance imaging multi-voxel pattern analysis. Eur. J. Neurosci. 42, 2851–2859 (2015).

33. J. Xiong, et al., Long-term motor training induced changes in regional cerebral blood flow in both task and resting states. NeuroImage 45, 75–82 (2009).

34. L. Ma, et al., Changes in regional activity are accompanied with changes in inter-regional connectivity during 4 weeks motor learning. Brain Res. 1318, 64–76 (2010).

35. V. B. Penhune, J. Doyon, Dynamic Cortical and Subcortical Networks in Learning and Delayed Recall of Timed Motor Sequences. J. Neurosci. 22, 1397–1406 (2002).

36. C. Horenstein, M. J. Lowe, K. A. Koenig, M. D. Phillips, Comparison of unilateral and bilateral complex finger tapping-related activation in premotor and primary motor cortex. Hum. Brain Mapp. 30, 1397–1412 (2009).

37. D. A. Nowak, C. Grefkes, M. Ameli, G. R. Fink, Interhemispheric competition after stroke: Brain stimulation to enhance recovery of function of the affected hand. Neurorehabil. Neural Repair 23, 641–656 (2009).

38. M. Kobayashi, H. Théoret, A. Pascual-Leone, Suppression of ipsilateral motor cortex facilitates motor skill learning. Eur. J. Neurosci. 29, 833–836 (2009).

39. B. W. Vines, D. G. Nair, G. Schlaug, Contralateral and ipsilateral motor e¡ects after transcranial direct current stimulation. NeuroReport 17, 672–674 (2006).

40. J. Duque, et al., Transcallosal inhibition in chronic subcortical stroke. NeuroImage 28, 940–946 (2005).

41. M. L. Harris-Love, E. Chan, A. W. Dromerick, L. G. Cohen, Neural Substrates of Motor Recovery in Severely Impaired Stroke Patients With Hand Paralysis. Neurorehabil. Neural Repair 30, 328–338 (2016).

42. N. Murase, J. Duque, R. Mazzocchio, L. G. Cohen, Influence of interhemispheric interactions on motor function in chronic stroke. Ann. Neurol. 55, 400–409 (2004).

43. A. Keitel, H. Øfsteng, V. Krause, B. Pollok, Anodal Transcranial Direct Current Stimulation (tDCS) Over the Right Primary Motor Cortex (M1) Impairs Implicit Motor Sequence Learning of the Ipsilateral Hand. Front. Hum. Neurosci. 12 (2018).

44. R. Hamel, et al., Bilateral intracortical inhibition during unilateral motor preparation and sequence learning. Brain Stimulat. 17, 349–361 (2024).

45. J. Diedrichsen, T. Wiestler, J. W. Krakauer, Two Distinct Ipsilateral Cortical Representations for Individuated Finger Movements. Cereb. Cortex 23, 1362–1377 (2013).

46. E. Berlot, G. Prichard, J. O’Reilly, N. Ejaz, J. Diedrichsen, Ipsilateral finger representations in the sensorimotor cortex are driven by active movement processes, not passive sensory input. J. Neurophysiol. 121, 418–426 (2019).

47. T. Wiestler, S. Waters-Metenier, J. Diedrichsen, Effector-independent motor sequence representations exist in extrinsic and intrinsic reference frames. J. Neurosci. Off. J. Soc. Neurosci. 34, 5054–5064 (2014).

48. S. Waters, T. Wiestler, J. Diedrichsen, Cooperation Not Competition: Bihemispheric tDCS and fMRI Show Role for Ipsilateral Hemisphere in Motor Learning. J. Neurosci. 37, 7500–7512 (2017).

